# Crystal structure of a novel guanine nucleotide exchange factor encoded by the scrub typhus pathogen *Orientia tsutsugamushi*

**DOI:** 10.1101/2020.06.04.133876

**Authors:** Christopher Lim, Jason M. Berk, Alyssa Blaise, Josie Bircher, Anthony J. Koleske, Mark Hochstrasser, Yong Xiong

**Affiliations:** Department of Molecular Biophysics & Biochemistry, Yale University, New Haven, CT 06520

**Keywords:** Scrub typhus, *Orientia*, Rho GTPases, Rac1, Cdc42, GEF, pathogen, effector

## Abstract

Rho family GTPases regulate an array of cellular processes and are often modulated by pathogens to promote infection. Here, we identify a cryptic guanine nucleotide exchange factor (GEF) domain in the OtDUB protein encoded by the pathogenic bacterium *Orientia tsutsugamushi*. A proteomics-based OtDUB interaction screen identified numerous potential host interactors, including the Rho-GTPases Rac1 and Cdc42. We discovered a new domain in OtDUB with Rac1/Cdc42 GEF activity (OtDUB_GEF_), with higher activity toward Rac1 *in vitro*. While this GEF bears no obvious sequence similarity to known GEFs, crystal structures of OtDUB_GEF_ alone (3.0 Å) and complexed with Rac1 (1.7 Å) reveal striking convergent evolution, with a distinct topology, on a V-shaped bacterial GEF fold shared with other bacterial GEF domains. Structure-guided mutational analyses identified residues critical for activity and a novel mechanism for nucleotide displacement. Ectopic expression of OtDUB activates Rac1 preferentially in cells, and expression of the OtDUB_GEF_ alone alters cell morphology. Cumulatively, this work reveals a novel bacterial GEF within the multifunctional OtDUB that co-opts host Rac1 signaling to evoke changes in cytoskeletal structure.

## Introduction

The Ras homologous (Rho) family of GTPases is part of the Ras superfamily of small G proteins. Rho family GTPases are molecular switches that control intracellular actin dynamics and regulate a diverse array of cellular processes from cytokinesis to cell migration and wound healing.^1-3^ These small ∼21 kDa proteins are highly conserved in all eukaryotes, with three founding family members that have been extensively studied: Rac1, Cdc42, and RhoA. Each Rho family GTPase exerts specific effects on the actin cytoskeleton, and constitutive activation of each protein leads to characteristic cellular phenotypes.

The signaling activity of a GTPase is controlled by its bound nucleotide state. When GDP is bound, the GTPase is in the “inactive” state, and loading of a GTP promotes the “active” conformation of the G protein. Interaction with downstream effector proteins and subsequent actin reorganization only occurs when the GTPase is in the GTP-bound “active” state. The intrinsic nucleotide exchange (GDP to GTP) and hydrolysis (GTP to GDP) rates of Rho family GTPases alone are slow. Nucleotide exchange occurs on the order of 1.5 per hour^4,5^, and the intrinsic hydrolysis rate is approximately 0.15 per min.^6^ Rapid regulation of GTPases, therefore, is controlled by two classes of proteins that either switch them “on” or “off”: <u>g</u>uanine nucleotide <u>e</u>xchange factors (GEFs) promote the dissociation of GDP and allow loading with GTP, and <u>G</u>TPase <u>a</u>ctivating <u>p</u>roteins (GAPs) accelerate the intrinsic GTP hydrolysis by the G protein.

Bacterial pathogens such as certain species of *Salmonella* and *Shigella* and enteropathic *E. coli*, encode and secrete effector proteins that modulate small GTPases to benefit the bacterium during infection. For instance, the *Shigella flexineri* IpgB2 GEF protein activates RhoA and causes characteristic membrane ruffles that are critical for *Shigella* invasion of the host cell.^7^ Bacterial effector GEFs belong to either the WxxxE family (named for a conserved motif important for folding and structural integrity) or the SopE family (SopE, SopE2, and BopE). These bacterial effectors share no sequence or structural homology to eukaryotic Rho GEFs, which predominantly belong to the Dbl homology (DH) family of GEFs typically formed by a six-helix bundle with an elongated, kinked “chaise lounge” fold.^8,9^ Rather, bacterial effector GEFs adopt a characteristic compact V-shaped fold, yet activate the Rho GTPases via the same contact regions in the GTPases that are crucial for nucleotide exchange by DH-family GEFs.^10^ While substantial effort has been exerted in detailing the molecular determinants of bacterial GEF activities and specificities, no bacterial effector GEFs have been identified outside of the WxxxE/SopE-like families.

Recently, we identified and characterized a putative effector protein, OtDUB, from the obligate intracellular bacterium that causes scrub typhus, *Orientia tsutsugamushi*. Despite extensive characterization of the OtDUB deubiquitylase (DUB) domain (residues 1-259), the function of the extensive C-terminal region, encompassing more than 1000 amino acids, remained elusive.^11^ Here, we report that the OtDUB encodes a novel GEF domain. Using biochemical, structural, and cellular methods, we demonstrate that the OtDUB_GEF_ predominantly activates Rac1 *in vitro* and *in vivo*, and that its three-dimensional structure differs drastically in topological and helical arrangement from both the WxxxE and SopE effector GEFs. We further determined the OtDUB_GEF_:Rac1 complex crystal structure, which reveals that despite its divergence in primary sequence and topology, the OtDUB_GEF_ has convergently evolved the V-shaped fold of bacterial GEFs and interacts with Rac1 at key common loci in the GTPase. The complex structure also suggested a distinct mechanism for GDP displacement, unique among all GEFs characterized to date. Our work reveals that *Orientia* has evolved a GEF domain that expands the molecular repertoire of bacterial effectors, and suggests a critical function for OtDUB in regulation regulating Rac1 to benefit the pathogen during infection.

## Results

### The OtDUB C-terminal segment is toxic in yeast and interacts with GTPases

The OtDUB N-terminal region contains an active DUB and a high-affinity ubiquitin binding domain (UBD) within the first 259 residues.^11^ However, the remainder of the 1369-residue protein is devoid of any computationally predicted domains. To examine how the OtDUB might affect eukaryotic cells, we first generated a series of truncations (Fig. 1a) and expressed the proteins in the yeast *Saccharomyces cerevisiae* (Fig. 1b). Remarkably, expression of the full-length protein caused a complete block to growth even if the DUB was inactivated by mutation of the catalytic cysteine, C135A (Figs. 1b and S1a). Expression of a fragment excluding the N-terminal DUB and UBD domains (OtDUB_275-1369_) also exhibited severe toxicity in yeast, but toxicity was no longer observed with the shorter OtDUB_675-1369_. The OtDUB_1-675_ truncation caused a lesser but still substantial growth deficit that was partially DUB dependent. These data suggest that a region within residues 275-1369 is the principal source of toxicity when expressed in yeast, but in the absence of this toxic domain, an additional DUB-dependent growth defect is uncovered.

**Figure 1.**
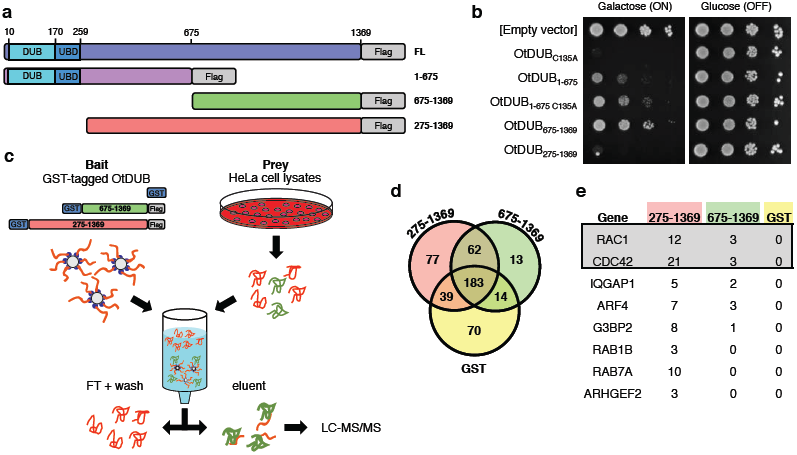
OtDUB_275-1369_ is toxic in yeast and binds multiple proteins in HeLa cell lysates. (a) Cartoon diagram of various OtDUB fragments used for ectopic expression in yeast, mammalian cell cultures, and bacteria. (b) Growth of W303 yeast expressing various OtDUB fragments from the galactose-inducible promoter in p416GAL1. Yeast cultures were serially diluted in ten-fold steps and spotted on SD-URA plates containing either galactose or glucose as the carbon source and grown for 3 days at 30°C. (c) Overview of LC-MS/MS experiment to identify host interactors of OtDUB fragments. Bait proteins immobilized on glutathione resin were incubated with HeLa cell lysates, washed and eluted by proteolytic cleavage from the GST tags using GST-human rhinovirus 3C protease (GST-HRV3C). (d) Venn diagram of total proteins identified between GST, OtDUB_275-1369_ and OtDUB_675-1369_. (e) Top candidate interactors of indicated OtDUB fragments from LC-MS/MS. Total peptide counts are shown.

To determine if specific human host proteins bind the OtDUB “toxic region,” we isolated proteins from HeLa cell lysates that specifically bound glutathione-S-transferease (GST)-tagged OtDUB fragments and identified them using liquid chromatography and tandem mass spectrometry (LC-MS/MS) (Fig. 1c). We identified several proteins that bound uniquely to GST-OtDUB_275-1369_ and not to GST-OtDUB_675-1369_ or the GST protein control (Fig. 1d, Supplementary Table 1). Notably, several small GTPases, such as Rac1 and Cdc42, and proteins of related functions were significantly enriched in the GST-OtDUB_275-1369_-bound sample compared to the GST-OtDUB_675-1369_ and GST controls (Fig. 1e). We focused on the small GTPases as there are numerous examples of pathogens hijacking GTPase signaling.^12,13^

**Table 1.**
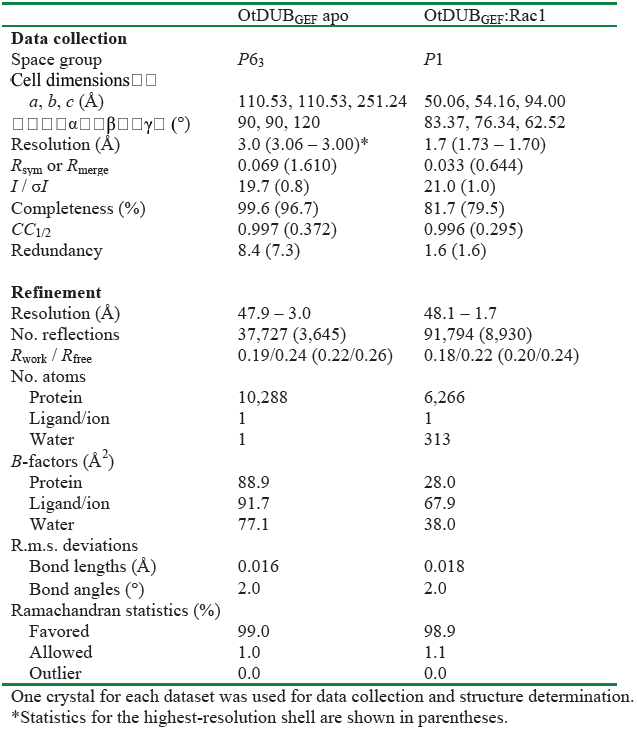
Data collection and refinement statistics.

### Identification of GTPase-binding and GEF activity in OtDUB

We carried out co-immunoprecipitation (co-IP) assays using lysates of HeLa cells ectopically expressing several OtDUB fragments to verify the putative interactions identified by mass spectrometry. Flag-tagged OtDUB fragments were immunoprecipitated following incubation of whole cell lysates, and bound proteins were resolved by SDS-PAGE and immunoblotted for Rac1, Cdc42, and RhoA. These co-IP experiments confirmed that full-length (FL) OtDUB and OtDUB_275-1369_ bound Rac1 and Cdc42 but not RhoA (Fig. 2a). Further truncations (OtDUB_1-675_ and OtDUB_675-1369_) abolished the interaction. To demonstrate that this association does not require other cellular components, we performed direct binding assays between purified *E. coli*-expressed recombinant OtDUB fragments and recombinant GST-tagged Rac1 or Cdc42. In agreement with the co-IP results with HeLa cell lysates, OtDUB_275-1369_-Flag bound to both Rac1 and Cdc42 in anti-Flag immunoprecipitation assays, whereas OtDUB_675-1369_-Flag did not. The reciprocal GST pulldowns produced analogous results (Fig. 2b).

**Figure 2.**
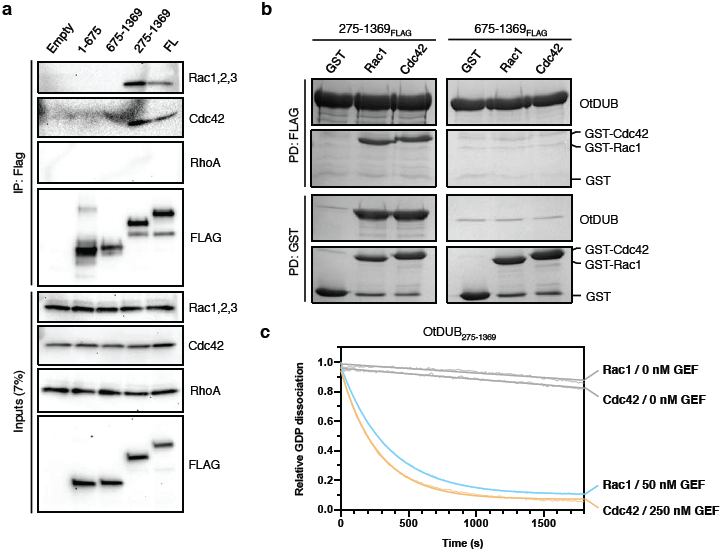
OtDUB binds Rac1 and Cdc42 and catalyzes nucleotide exchange *in vitro*. (a) Inputs and anti-Flag immunoprecipitates of lysates from HeLa cell ectopically expressing the indicated Flag-tagged OtDUB fragments. Proteins were resolved by SDS-PAGE and immunoblotted for Rho GTPases (Rac1,2,3; Cdc42; RhoA). (b) FLAG (top) and reciprocal GST (bottom) pulldown experiments between purified recombinant FLAG-tagged OtDUB fragments and GST-tagged Rac1 or Cdc42. Proteins were resolved by SDS-PAGE and stained with Coomassie Blue. (c) Time course of the dissociation of BODIPY-GDP from Rac1 or Cdc42 (9 μM) in the presence of OtDUB_275-1369_ (0, 50, or 250 nM) as measured by loss of BODIPY-GDP fluorescence. Excitation and emission wavelength were 488 nm and 535 nm, respectively.

GTPase-activating proteins (GAPs) and guanine nucleotide exchange factors (GEFs) are the most common binding partners for GTPases. Several observations suggested that the OtDUB fragment might harbor GEF activity. First, certain intracellular bacterial pathogens encode GEFs to subvert GTPase signaling and alter actin networks during infection.^14^ Additionally, the preceding pulldowns required EDTA, a condition that promotes nucleotide unloading and enhances GEF:GTPase interactions.^15^

To test whether OtDUB has GEF activity, we measured the rate of dissociation of a fluorescent GDP analog (BODIPY-GDP) from Rac1 and Cdc42.^16^ BODIPY-GDP only fluoresces strongly when bound to protein. In the absence of OtDUB, Rac1 and Cdc42 released very little BODIPY-GDP (Fig. 2c). In contrast, OtDUB_275-1369_ (50 nM) greatly accelerated nucleotide dissociation from Rac1. The same fragment also promoted GDP release from Cdc42, albeit less efficiently, requiring five times more OtDUB (250 nM) to achieve similar levels of dissociation. In line with our initial binding studies, OtDUB_275-1369_ did not catalyze GDP dissociation from RhoA, and the shorter OtDUB_675-1369_ fragment displayed no exchange activity with Rac1 (Fig. S1b). Together, these data indicate that OtDUB_275-1369_ binds to and promotes nucleotide exchange on both Rac1 and Cdc42.

### Determination of a minimal, GTPase-binding OtDUB_GEF_ domain

Bacterial GEFs in the WxxxE/SopE family exclusively target Rho GTPases yet are divergent in both sequence and structure from all eukaryotic Rho GEFs. Intriguingly, no region of the OtDUB protein sequence aligns to WxxxE/SopE sequences nor to any eukaryotic GEF sequence. Hence, we employed a truncation mapping strategy to further define the boundaries of a potential GEF domain in OtDUB. Guided by secondary structure predictions, we generated nine OtDUB constructs and tested them for Rac1 binding in an *in vitro* GST pulldown assay (Fig. 3a). Consistent with our co-IP results, the ∼1000-residue (OtDUB_275-1369_) fragment strongly associated with Rac1. Neither C-terminal truncation (275-913) nor N-terminal truncations up to residue 548 reduced binding to Rac1; however, N-terminal truncation to residue 579 abolished binding. Further truncation from the C-terminus revealed a minimal, ∼200-residue fragment (residues 548-759, hereafter referred to as OtDUB_GEF_) that retained full binding to Rac1.

**Figure 3.**
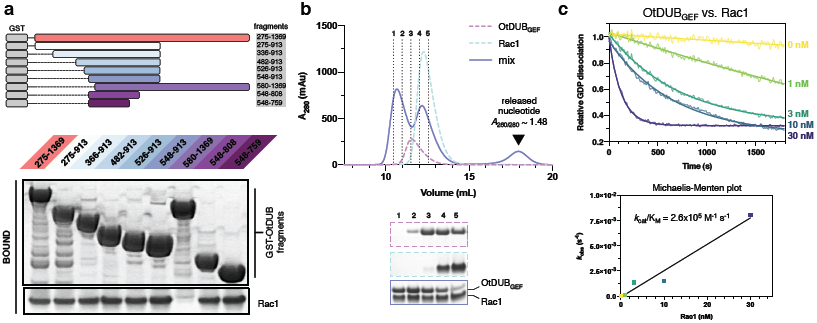
A 200-residue OtDUB subdomain sufficient for binding Rac1. (a) OtDUB truncations used for mapping studies (top) and GST pulldown experiment using GST-OtDUB fragments as bait and Rac1 as prey (bottom). Gel was stained with Coomassie Blue. N-terminal truncation to residue 580 abolished binding, whereas the 548-759 fragment (OtDUB_GEF_ domain) retained full binding capacity. (b) Size exclusion chromatography of OtDUB_GEF_:Rac1 mixtures demonstrates stable complex formation (top). Column fractions were evaluated by SDS-PAGE and protein staining (bottom). (c) Time course of the dissociation of BODIPY-GDP from Rac1 in the presence of increasing amounts of OtDUB_GEF_ (residues 548-759). Raw fluorescence curves were fit to a single exponential decay (top), and initial rates were plotted against Rac1 concentration (bottom). Linear transformation of the titration yielded a *k*_cat_/K_M_ of 2.6 ± 0.3 ×10^5^ M^-1^ s^-1^.

We employed size exclusion chromatography (SEC) to confirm that OtDUB_GEF_ behaved well in solution and formed a stable complex with Rac1. Indeed, free OtDUB_GEF_ eluted as a monodisperse peak, and its mixture with Rac1 resulted in co-elution of both proteins at the expected volume for a 1:1 complex (Fig. 3b). Furthermore, another new peak appeared in the mixture fractionation at an elution volume corresponding to that of a small molecule; the high *A*_260/280_ ratio of this peak (∼1.5) suggested it represents released nucleotide that had co-purified with Rac1. Similar SEC profiles were obtained for Cdc42 (Figs. S2a and S2b), demonstrating stable complex formation. We used isothermal titration calorimetry (ITC) to quantify the binding affinity of OtDUB_GEF_ toward Rac1 or Cdc42 (bound to co-purified nucleotides). The dissociation constant (*K*_d_) between OtDUB_GEF_ and Rac1 is approximately 5 μM, similar to those of other GEF:GTPase pairs (Fig. S2c).^17-19^ The measured *K*_d_ for Cdc42 is ∼16 µM. This three-fold difference in affinity may partially explain the weaker activation of Cdc42 by OtDUB_GEF_.

Finally, to ensure that the OtDUB_GEF_ fragment fully accounts for catalytic activity, we tested various concentrations of this fragment for GEF activity against Rac1. In agreement with results from the pulldown assay, OtDUB_GEF_ catalyzed GDP dissociation from Rac1 at all concentrations tested (Fig. 3c). Extraction of initial dissociation rates from this titration resulted in a catalytic efficiency of *k*_cat_/K_M_ = 2.6 × 10^5^ M^-1^ s^-1^, comparable in magnitude to SopE acting on Rac1 (5.0 × 10^5^ M^-1^ s^-1^).^20^. In contrast, OtDUB_GEF_ was a relatively poor GEF against Cdc42, with a catalytic efficiency approximately 15-fold worse than that for Rac1 (Figs. S2d and S2e). Together, our *in vitro* findings demonstrate that the ∼200-residue OtDUB_GEF_ fragment is sufficient to promote nucleotide exchange in two different Rho GTPases. The remainder of our studies focused solely on Rac1 because of the clear preference of OtDUB_GEF_ for this GTPase.

### Crystal structure of OtDUB_GEF_

All prokaryotic GEF effectors with available structures diverge from the most common eukaryotic Rho-GEF family, the Dbl-homology (DH) GEFs, which typically adopt an extended helical bundle, or “chaise lounge” fold.^9^ We crystallized OtDUB_GEF_ (residues 548-759 of OtDUB) in its apo form and determined the structure to 3.0 Å resolution. The structure reveals that OtDUB_GEF_ is composed exclusively of alpha helices and random coil regions (Figs. 4a and S5a), and adopts the familiar V-shaped fold seen in WxxxE/SopE GEFs (Fig. 4b). Despite their structural similarity at the level of the overall protein fold, the topology of OtDUB_GEF_ is very distinct. One conserved feature of all previously resolved bacterial effector GEFs is their shared “back-and-forth” topology of helices.^14^ In SopE and Map, for example, α1/4/5 form the respective “left” lobes, and α2/3/6 form the “right” lobes of both proteins (Fig. 4c). By contrast, OtDUB_GEF_ splits sequentially into an N-terminal lobe (α1-3) and a C-terminal lobe (α4-9) (Figs. 4a and 4c). The lack of similarity in primary sequence renders alignment impossible in the case of OtDUB_GEF_ and precludes prediction from amino acid sequence. The striking divergence of topology and sequence for a common fold suggests convergent evolution between the OtDUB_GEF_ and SopE/WxxE family GEFs.

**Figure 4.**
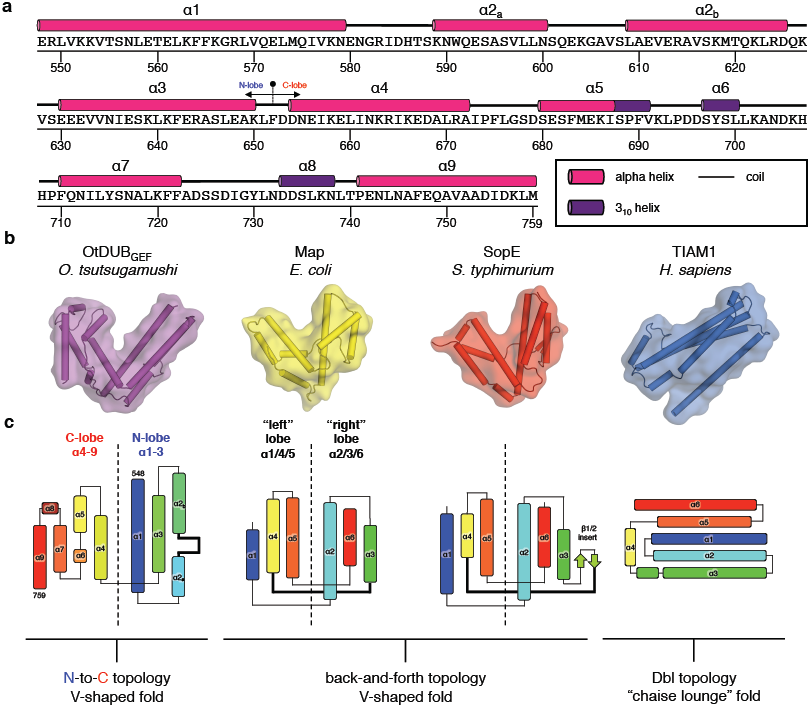
Crystal structure of the apo OtDUB_GEF_ domain reveals a novel topology. (a) Secondary structure diagram of OtDUB_GEF_. Alpha and 3_10_ helices are represented as cylinders in magenta and purple, respectively, and random coils are represented as solid lines. (b) Overall structural comparison of apo OtDUB_GEF_ and other Rho GEFs: Map (PDB: 3GCG), SopE (PDB: 1GZS), TIAM1 (PDB: 1FOE). All structures shown with helices as cylinders and transparent surface. (c) Topology diagrams of the same GEFs color-ramped from N-terminus (blue) to C-terminus (red) to highlight the unique topology of OtDUB_GEF_. Helices are shown as rectangles, beta strands are shown as arrows, and loop regions are shown as lines.

### Crystal structure of the OtDUB_GEF_:Rac1 complex

To gain insight into the mechanism of Rac1 activation by OtDUB_GEF_, we determined a crystal structure of the OtDUB_GEF_:Rac1 complex to 1.7 Å resolution (Fig. 5A). OtDUB_GEF_ does not undergo major conformational changes upon binding to Rac1, with overall RMSDs ranging from 0.1Å to 1.9 Å when compared to the six independent copies of the molecule found in the asymmetric unit of the apo crystal (Fig. S5b), suggesting a rigid structure evolved to be primed for interaction. Like other Rho-GEFs, OtDUB_GEF_ forms contacts with three key surfaces of the Rac1 GTPase: the switch I and II regions, residues near the nucleotide-binding cleft (the catalytic pocket), and the β2-3 hairpin “interswitch” region, burying a total surface area of more than 1,800 Å^2^ per molecule (Fig. 5a). Each interacting region is described in detail below.

**Figure 5.**
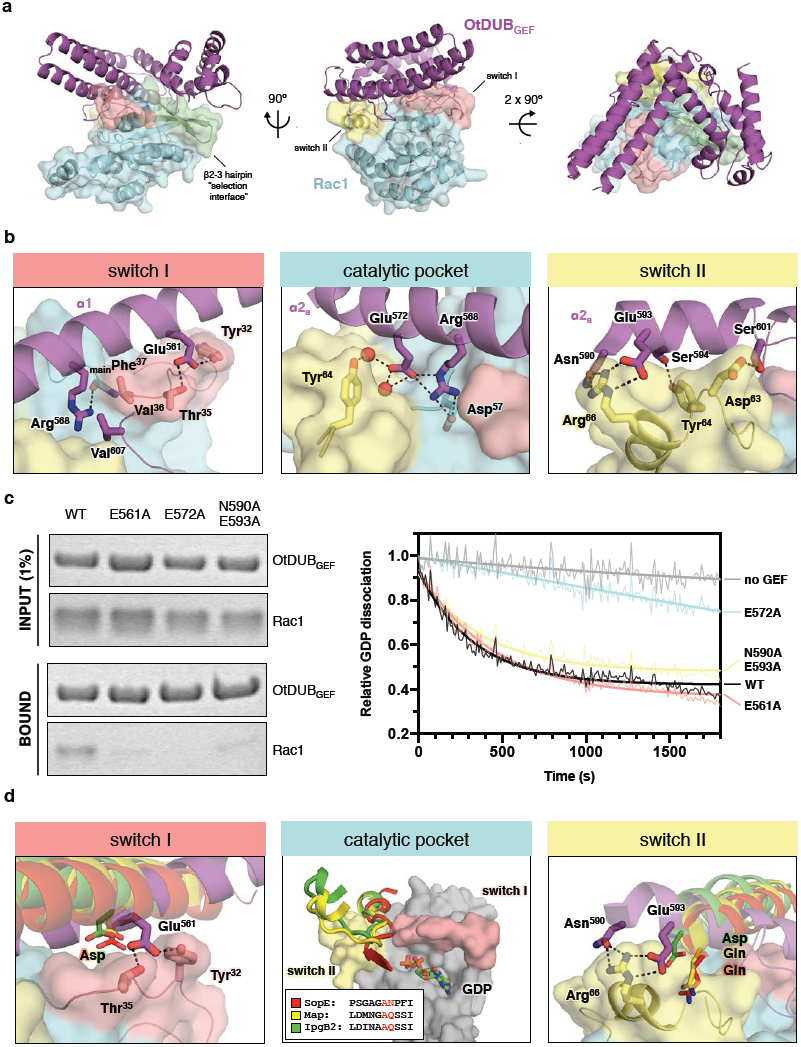
Biochemical function and structure of the OtDUB_GEF_:Rac1 complex. (a) Three orthogonal views of the complex with OtDUB_GEF_ in purple and Rac1 in cyan. Switch I and II loops are in salmon and yellow, respectively, and the β2-3 hairpin selectivity interface is green. (b) Close-up views of each key interface between OtDUB_GEF_ and Rac1 showing selected residues, with hydrogen bonds and electrostatic interactions shown as dashes, water molecules as red spheres. (c) GST pulldown assays (left) of GST-OtDUB_GEF_ (WT or charge-neutralizing mutations at each interface) incubated with Rac1 and analyzed by SDS-PAGE and Coomassie Blue staining. Corresponding BODIPY-GDP release assays are shown on the right. (d) Close-up views of each interface comparing the OtDUB_GEF_ residues interacting with Rac1 (thick sticks) to the interactions made by other bacterial GEFs (thin sticks).

The switch I interface (salmon in Fig. 5b, left) is composed of a hydrophobic pocket formed by OtDUB_GEF_-Phe^564^/Val^607^/Val^612^ that accommodates Rac1-Val^36^. This hydrophobic cleft is flanked on one side by OtDUB_GEF_-Arg^568^ interacting with the main chain carbonyl of Rac1-Phe^37^, and on the other by OtDUB_GEF_-Glu^561^ forming a polar network with Rac1-Thr^35^ and Tyr^32^. At the switch II interface (yellow in Fig. 5b, right), OtDUB_GEF_ forms extensive electrostatic interactions, coordinating Rac1-Arg^66^ via OtDUB_GEF_-Asn^590^ and Glu^593^. Furthermore, OtDUB_GEF_ residues Ser^594^, Ser^601^, and Glu^603^ form charged-polar interactions with Rac1 residues Tyr^64^, Asp^63^, and the backbone of Glu^62^, respectively. Finally, near the nucleotide-binding pocket, an extensive network of ordered water molecules is coordinated by OtDUB_GEF_-Glu^572^, which interact with multiple Rac1 residues (Fig. 5b, center).

To confirm the importance of these residues, we mutated important OtDUB_GEF_ residues at each interface and tested these mutants for both binding and catalytic activities against Rac1. Compared to wild-type OtDUB_GEF_, both switch I (E561A) and switch II (N590A/E593A) mutants reduced apparent binding in the GST pulldown assay, whereas little defect in catalysis was observed (Fig. 4c). By contrast, mutation of the central catalytic residue (E572A) abrogated binding entirely and reduced GDP dissociation activity to background levels, highlighting the importance of this central residue for interaction with Rac1. This mutational analysis confirms that OtDUB harbors a *bona fide* GEF domain with a unique topology.

The OtDUB_GEF_:Rac1 complex further revealed the structural basis for its selectivity amongst Rho-family GTPases. Previously, a “lock-and-key” mechanism for substrate selection has been proposed for discrimination between Rac1, Cdc42, and RhoA.^21,22^ In this model, complementary pairing of amino acids generates favorable interactions for target GTPases in the β2-3 hairpin “selectivity patch”, and steric hindrance precludes interaction with non-target family members. For instance, where Rac1 and Cdc42 encode small amino acids, Ala^3^ and Thr^3^, respectively, RhoA places a bulky, charged Arg^5^ residue at the same position (Fig. S3a). In agreement with the dual specificity observed in the *in vitro* GEF assay, OtDUB_GEF_ residues are compatible with both selectivity patches of Rac1 and Cdc42. In contrast, the bulky, charged amino acids of RhoA (Arg^5^ and Glu^54^) are predicted to clash with several rigid regions of OtDUB_GEF_ (Fig. S3b).

We next analyzed whether OtDUB_GEF_ makes similar contacts with Rac1 when compared other bacterial GEFs and their cognate Rho-family GTPases: SopE/Cdc42 (PDB: 1GZS), Map/Cdc42 (PDB: 3GCG), and IpgB2/RhoA (PDB: 3LW8, complex A). Overall, the footprints of the bacterial effector GEFs and OtDUB_GEF_ on their GTPases are very similar (Fig. S4). However, comparison of the interactions made by WxxxE/SopE GEFs (hereafter referred to collectively as bacGEFs) with OtDUB_GEF_ revealed that the latter has distinct contacts with Rac1. Interactions with switch I are the most similar: where bacGEFs use an invariant acidic Asp residue to make backbone contacts with Val^36^ and Thr^38^ (Rac1/Cdc42 numbering), OtDUB_GEF_ positions Glu^561^ to interact with Rac1-Tyr^32^ and Thr^35^ (Fig. 5d, left). Switch II interactions are slightly more diverse in bacGEFs, with SopE and Map using a Gln residue to make polar contacts with Asp^65^, and IpgB2 positioning an Asp sidechain to interact with Arg^66^ of RhoA. Here, OtDUB_GEF_ is also unique, coordinating Rac1-Arg^66^ with Asn^590^/Glu^593^, and interacting with Tyr^64^ instead of Asp^65^ (Fig. 5d, right). Finally, WxxxE/SopE GEFs extrude flexible “catalytic loops” between α3 and α4 that rest in a cleft formed between switch I and switch II of the GTPase (Fig. 5d, center). These catalytic loops share some sequence similarity, requiring an Ala followed by a polar Asn/Gln residue to facilitate nucleotide exchange. Strikingly, the unique topology of OtDUB_GEF_ necessarily precludes the existence of a homologous catalytic loop, suggesting that OtDUB_GEF_ utilizes a novel catalytic mechanism.

### OtDUBGEF utilizes an innovative carbonyl “catalytic lever” to dissociate nucleotide

In all DH GEF:Rho GTPase structures, highly conserved and characteristic conformational changes in switch II are responsible for dissociation of GDP. For instance, the canonical GEF-bound structure of the TIAM1:Rac1 complex (PDB: 1FOE, slate) reveals that the sidechains of Glu^62^ and Ala^59^ of Rac1 both rotate nearly 180° toward the nucleotide-binding cleft relative to their positions before GEF binding, i.e. in the GDP-bound state (PDB: 5N6O, grey) (Fig. 6a). This allows Rac1-Glu^62^ to interact with Rac1-Lys^16^, competing for γ-phosphate binding, and Rac1-Ala^59^ displaces Mg^2+^ in a hydrophobic pocket (Fig. 6a). Near identical conformations are observed in all other bacterial GEF complex structures for SopE:Cdc42 and Map:Cdc42, and interactions with their catalytic loops facilitate this conformational change.^23,24^ Thus, all known Rho GEFs use a similar catalytic mechanism that drives the GTPase switch I region through a conserved set of main chain and side chain motions toward the bound GDP. Accordingly, mutation of Rac1-Glu^62^ abrogates dissociation activity for all Rho GEFs tested to date, as well as for most Ras and Ran GEFs.^25^

**Figure 6.**
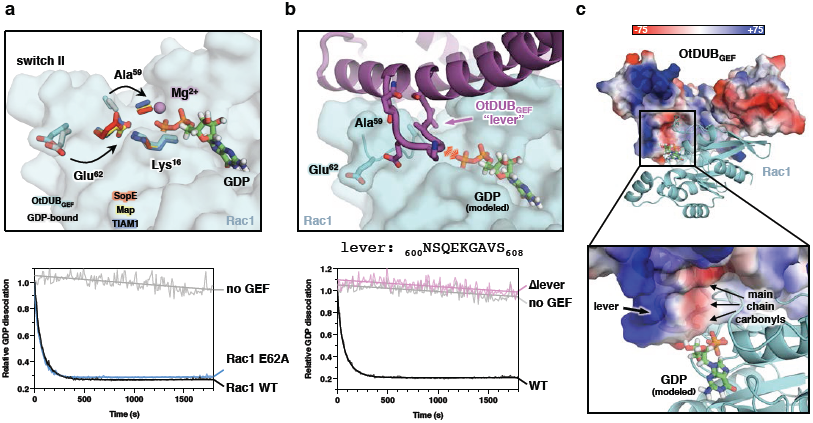
OtDUB_GEF_ utilizes a unique carbonyl “catalytic lever” to catalyze nucleotide exchange. (a) Top: Structural overlay of GTPases from GDP-bound (PDB: 5N6O, grey) and GEF-bound structures: OtDUB_GEF_ (cyan), SopE (red), Map (yellow), and TIAM1 (slate). GEFs are removed for clarity. Switch I residues Glu^62^, Ala^59^, and Lys^16^ are shown as sticks. Arrows indicate the previously observed motions of the residues upon GEF binding. Bottom: BODIPY-GDP exchange assay reveals that Glu^62^ is not required for efficient exchange catalyzed by OtDUB_GEF_. (b) Top: A loop of OtDUB_GEF_ that interrupts α2 acts as a “catalytic lever” to promote release of GDP. OtDUB_GEF_ is shown as cartoon (purple), and Rac1 is shown as transparent surface (cyan). GDP (not present in our structure) is modeled based on PDB 5N6O for reference. The steric clash and electrostatic repulsion are indicated by the red lines. Bottom: Nucleotide exchange assay reveals a strict requirement in the lever segment for exchange of BODIPY-GDP. (c) Calculated electrostatic surface potential map (unit k_B_T/e) shows high negative charge in the OtDUB_GEF_ lever formed by a tri-carbonyl motif at the vicinity of the diphosphate group of GDP.

Strikingly, the conformation of Rac1-Glu^62^ in our OtDUB_GEF_:Rac1 complex is swung away from the nucleotide pocket, and instead resembles the GDP-bound conformation (Fig. 6a). Our structure was obtained after nucleotide dissociation (Fig. 3c) and clearly show no nucleotide present (Fig. S5b), suggesting that we captured the complex in a post-exchange conformation despite the observed pre-exchange position of Rac1-Glu^62^. This discrepancy led us to hypothesize that Rac1-Glu^62^ would be dispensable for GDP dissociation. Indeed, OtDUB_GEF_-catalyzed nucleotide dissociation was indistinguishable between wild-type and E62A Rac1 (Fig. 6a). There are exceptions to the dependence on Glu^62^ (or its equivalent), as in the case of the GEF domain of Rabex-5 and its GTPase Rab21, where Rabex-5 supplies an acidic residue *in trans* to catalyze exchange on Rab21. ^26^ However, to our knowledge OtDUB_GEF_ is the first example of a Rho-specific GEF whose catalytic mechanism does not rely on Glu^62^.

How does OtDUB_GEF_ promote nucleotide exchange in the absence of a canonical catalytic loop? In addition to the unusual conformation of Glu^62^ in our complex structure, we also observed a short loop in OtDUB_GEF_ (_601_SQEKGAVS_608_, hereafter referred to as the “catalytic lever”) that protrudes out of α2 near the canonical catalytic loop position (Fig. 6b). The deep projection of the OtDUB_GEF_ lever into the nucleotide-binding cleft led us to hypothesize that this loop may be critical for nucleotide exchange. Deletion of the catalytic lever in OtDUB_GEF_ completely abrogated nucleotide exchange activity compared to wild-type *in vitro* (Fig. 6b). The compromised catalytic activity cannot be attributed to misfolding of the protein or to loss of binding Rac1 since the OtDUB_GEF_ Δlever protein was monodisperse and continued to form a complex with Rac1 (Fig. S5c). Inspection of the electrostatic surface potential of the catalytic lever of OtDUB_GEF_ near the nucleotide-binding cleft of Rac1 revealed significant negative charge formed not by side chains, but by three main chain carbonyls pointed towards the nucleotide-binding pocket (Fig. 6c). We propose that these polar carbonyl moieties repulse the negatively charged phosphates of GDP and thereby facilitate nucleotide exchange. The conformations of the catalytic lever in the OtDUB_GEF_ apo and Rac1-complexed crystal structures are remarkably similar (Fig. S5d, pairwise Cα RMSD_lever_ = 0.04 – 0.25 Å when comparing the two independent copies of the levers in the complex crystal with the six independent copies in the apo crystal), suggesting the catalytic lever is rigid and primed for direct interaction with Rac1. The attribution of catalysis activity to the catalytic lever highlights a unique mechanism utilized by OtDUB_GEF_ to promote nucleotide dissociation.

### OtDUB_GEF_ specifically activates Rac1 in cells

To complement these structural and *in vitro* activity results, we examined the *in vivo* activity state of Rac1 and Cdc42 in HeLa cells ectopically expressing OtDUB_FL_ or the GEF impaired mutant OtDUB_E572A_. We utilized a resin-bound GST fusion of the p21-binding domain (PBD) of Pak1 to specifically enrich the active forms of Cdc42 and Rac1.^27^ OtDUB-expressing cells exhibited a 2.7 ± 0.3 fold increase in activated Rac1 compared to control cells (Fig. 7a). Importantly, the E572A mutant, which abolishes OtDUB_GEF_ activity towards Rac1 *in vitro*, no longer raised active Rac1 levels in cells above baseline levels. Consistent with the lower *in vitro* activity of OtDUB_GEF_ on Cdc42 (Figs. 2c and S1e), there was no significant increase in the active Cdc42 when the OtDUB was ectopically expressed (1.1 ± 0.2 fold increase). These data both validate that the Rho-GTPase GEF activity is contained within the OtDUB_GEF_ domain and show a clear preference for Rac1 over Cdc42 *in vivo*.

**Figure 7.**
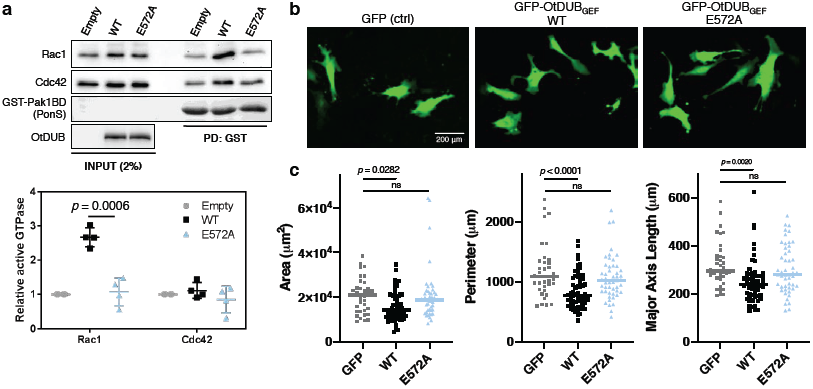
OtDUB_GEF_ specifically activates Rac1 and modulates cell morphology. (a) Lysates from HeLa cells carrying empty vector or plasmids expressing wild-type or E572A OtDUB were subjected to GST-Pak1PBD pulldowns to enrich active Rac1 and Cdc42. Representative experiment (left) from 4 independent experiments that were quantified (right) relative to input levels and the empty vector control. Bars represent mean and S.D. *P* value determined from a two-tailed unpaired Student t-test. Outlier from Rac1 wild-type (5.4 fold increase in activity) excluded one trial. Outlier was identified using Grubbs algorithm with alpha = 0.05. (b) Representative epifluorescence images of fibroblasts carrying the vector expressing only GFP or plasmids expressing wild-type or E572A OtDUB_GEF_. (c) Quantification of cell area, perimeter, and major axis length.

To examine possible downstream effects of OtDUB_GEF_-activated Rac1 on the actin cytoskeleton, we turned to a model cell line, mouse 3T3 fibroblasts. Ectopic expression of full-length OtDUB-GFP (WT or E572A) caused massive cell death and GFP signal could not be detected (data not shown). In contrast, expression of a GFP fusion of the GEF domain alone was tolerated by fibroblasts and expression was readily detected (Fig. 7b). Expression of OtDUB_GEF_ did not globally modulate filamentous actin structures, as revealed by phalloidin staining and confocal microscopy (Fig. S6).

Expression of OtDUB_GEF_ leads to statistically significant cell morphology changes. Rac1 regulation of the actin cytoskeleton promotes cell edge protrusion and Rac1 activity is required for cell spreading on adhesive surfaces in many cell types.^28-30^ Cells expressing the wild-type OtDUB_GEF_-GEF were slightly smaller, less elongated, and had fewer filopodial extensions compared to control cells expressing GFP alone (Fig. 7b). Quantification of cell shape metrics revealed that overall cell area, cell perimeter, and major axis length decreased upon expression of the OtDUB_GEF_-GFP (Fig. 7c). Critically, cells expressing the catalytically inactive OtDUB_GEF_-E572A-GFP mutant were not different from GFP control. Together, these data suggest that OtDUB_GEF_ activity is responsible for the observed morphological phenotypes, and that its GEF activity potentially regulates the host cytoskeletal machinery to produce distinct morphological defects.

## Discussion

The results presented here uncover an unexpected GEF activity in the multidomain OtDUB protein from the pathogen *Orientia tsutsugamushi* and provide unexpected insight into our understanding of the regulation of Rho GTPases. The cryptic GEF domain harbors GEF activity toward both Rac1 and Cdc42, with a clear preference for Rac1 both *in vitro* and in living cells. High-resolution crystal structures of the OtDUB_GEF_ domain alone and in complex with Rac1 reveal a new topology amongst reported GEFs that nevertheless adopts a conserved V-shaped GEF fold. Importantly, OtDUB_GEF_ utilizes a novel mechanism to activate Rho GTPases involving direct, steric exclusion of GDP and distinct molecular determinants.

The unique topology of OtDUB_GEF_ compared to all other bacterial effector GEFs represents a unique solution to a common objective for bacterial intracellular pathogens, namely, the manipulation of host GTPases to promote the survival and proliferation of the pathogen. Sequence comparison of the OtDUB_GEF_ domain against the nonredundant NCBI protein database revealed clear conservation only within the *Orientia* genus. Interestingly, a duplicated segment (residues 289-499) within the OtDUB protein itself shares 48% identity with the OtDUB_GEF_ domain (residues 548-759; Fig. S7a). This duplicated region does not have Rho-GTPase GEF activity nor does it bind Rho-GTPases (Fig. 2a); however, we cannot rule out that this region does not bind other small GTPases identified in our proteomics screen (Fig. 1e). Furthermore, the ortholog from *O. chuto* (OcDUB, WP_052694629.1) contains three OtDUB_GEF_ duplications: residues 429-641 with 47% identity to OtDUB_GEF_, 668-883(37% identity), and 928-1137 (50% identity) (Fig. S7b).

To our knowledge, the nucleotide exchange mechanism employed by OtDUB_GEF_ is unique amongst known GEFs. The role of Glu^62^ (or its equivalent) has been studied extensively by Gasper *et al*; whereas DH/Dbl Rho-GEFs (e.g. Dbs, TIAM1, and p190) and Cdc25 domain-containing Ras-GEFs (e.g. Sos and Rap) are absolutely dependent on Glu^62^ for nucleotide exchange, Sec7-like Arf-GEFs (e.g. Rabex-5 and Sec7) circumvent this requirement by supplying an acidic residue *in trans*.^26,31^ Notably, OtDUB_GEF_ exchange activity is independent of Glu^62^, and the structure does not implicate any nearby acidic residue from the GEF for interaction with Lys^16^, in contrast to Sec7-like GEFs. Instead, the catalytic lever of OtDUB_GEF_ inserts directly into the nucleotide pocket, revealing an unusual steric mechanism to promote GDP unloading. From the level of protein topology to the mechanistic details of nucleotide exchange, OtDUB_GEF_ is a fascinating example of convergent evolution, and represents a unique solution for interaction with a highly conserved master regulator of the host cytoskeleton, Rac1.

While it remains to be demonstrated that OtDUB is in fact a *bona fide* effector protein, the robust Rac1 activation in cultured cells further bolsters this possibility. The combination of the DUB, UBD, and GEF activities—all of which specifically interact with eukaryotic cellular signaling molecules that are absent from prokaryotic systems—strongly suggests that OtDUB is exposed to the host cellular milieu. We note that ectopic expression of OtDUB with inactivated DUB, UBD and GEF domains (OtDUB-C135A,V203D,E572A) still causes complete growth inhibition in yeast, suggesting additional regions of unknown function in OtDUB that can also interfere with host physiology (Fig. S8).

Due to a lack of early Rac1 activation during *Orientia* infection^32^, it is unlikely that the OtDUB_GEF_ acts in the same fashion as the SopE and SopE2 GEFs from *Salmonella*, which are secreted into the host cell cytoplasm to activate Rac1/Cdc42, inducing membrane ruffling and facilitating uptake of the bacterium.^33^ Rather, it is tempting to speculate that the OtDUB_GEF_ activates Rac1 after the bacterium escapes the vacuolar compartment and accesses the cytoplasmic space, roughly 2 hours post infection.^34^ One potential indicator of Rac1 activation during post-endosomal escape is the activation of the MAPK signaling pathway^35^, a known downstream target of Rac1.^36-38^ Additionally, IQGAP1, a downstream effector of Rac1/Cdc42 was found enriched in the OtDUB_275-1369_ mass spectrometry analysis. IQGAP1 is a scaffolding protein with numerous binding partners and roles^39^ and is also required for efficient infection by *Salmonella*.^40^

Cumulatively, the biochemical, bioinformatic, and structural data presented here demonstrate that a recently evolved GEF domain with a unique sequence, topology, and nucleotide-exchnge mechanism exists in the *Orientia tsutsugamushi* scrub typhus pathogen. The characterization of nearly half a dozen distinct and evolutionarily unrelated GEF families in eukaryotes indicates that numerous protein folds can support guanine nucleotide exchange activity. The unique topology of OtDUB_GEF_ further suggests that the sequence space of GEFs may be incompletely characterized. Finally, our work suggests that OtDUB GEF activity may be crucial for *Orientia* infection and opens the door for future investigations in this neglected pathogen.

## Methods

### DNA and cloning

The open reading frame (ORF) of *OTT_1962* from *Orientia tsutsugamushi* (Ikeda isolate) was synthesized (by Genscript) with codons optimized for human (primary) and yeast (secondary) expression and inserted into pCDNA3.1(+).^11^ The synthesis included a C-terminal Gly-Ser-Gly-1xFlag epitope sequence and 5’ BamHI, 3’ XhoI restriction sites. This DNA sequence was used as template for all the OtDUB plasmids in this study. All yeast and mammalian expression plasmids were generated by PCR amplification from the *OTT_1962* ORF with flanking BamHI/XhoI sites and where applicable included sequence for a C-terminal flag tag as above. All OtDUB fragments were cloned into pGEX6P1 (GE Healthcare) using BamHI/XhoI sites and included a sequence encoding a C-terminal Flag tag as described above. Coding sequences for Rac1, Cdc42, and RhoA were cloned into the BamHI/Not1 sites of pGEX6P1. DNA encoding residues 1-177 of both human *RAC1* and *CDC42* were cloned into an unmodified pET-28a (Addgene) vector that encodes an N-terminal TEV-cleavable 6xHis-tag using the NdeI/XhoI sites. For OtDUB, a sequence encoding residues 548 to 759 from the *OTT_1962* ORF were cloned similarly. For crystallization constructs, an uncleavable, C-terminal 6xHis-tag was introduced into both the OtGEF fragment and Rac1. Rac1 and Cdc42 template DNA was kindly provided by Titus Boggon. Standard site-directed mutagenesis was carried out for generating mutations. All PCR-generated plasmid inserts were verified by DNA sequencing.

### Yeast culturing and spot assays

Yeast were grown under standard conditions at 30°C using yeast rich (yeast extract-peptone-dextrose or YPD) or minimal (SD) media.^41^ Isolated transformants carrying p416GAL1-based plasmids^42^ in the W303 (MHY2416)^43^ *S. cerevisiae* background were grown overnight in SD-URA medium supplemented with casamino acids. Cultures were diluted in sterile water to 0.2 OD_600_ from which 10-fold dilutions were made. Diluted cultures were spotted onto SD-URA plates supplemented with either 2% glucose or 2% galactose. Plates were incubated at the indicated temperature for 3 days. Steady state levels of each OtDUB polypeptide was verified by back diluting 2.5 OD units of SD-URA + 2% raffinose cultures into SD-URA + 2% galactose, growing for 4h at 30°C and processing 2.5 OD units by alkaline lysis for subsequent Western immunoblot analysis using anti-Flag antibodies.

### Human cell culturing and transfections

HeLa cells (ATCC) were cultured in DMEM supplemented with 10% (v/v) fetal bovine serum and 1% (v/v) penicillin/streptomycin) in a 37° C humidified incubator containing 5% CO_2_. For transient transfections, 2.2×10^6^ HeLa cells were plated on a 10-cm cell culture dish, and transfected 24 h later using Xtreme-GENE 9 (Roche) according to the manufacturer’s directions. In brief, 750 μl Opti-MEM, 30 μl transfection reagent and 10 μg DNA were incubated together for 15 min before being added drop-wise to plated HeLa cells in antibiotic-free medium. Cells were collected 24 to 48 h post-transfection by scraping in ice-cold phosphate-buffered saline (PBS) and were pelleted and either processed immediately or flash frozen in liquid N_2_ and stored at −80°C.

Mouse 3T3 (WT32) fibroblast cells were maintained in DMEM plus 10% fetal bovine serum (FBS), 2 mM L-glutamine, and penicillin/streptomycin at 37° C and 5% CO_2_. 2.6×10^5^ cells were plated on a 10-cm dish the day before transfection and were transfected with plasmids pEGFP-N (expressing GFP alone), pEGFP-N-OtDUB-GFP, pEGFP-N-OtDUB-E572A-GFP, pEGFP-N-OtDUB_GEF_-GFP (residues 548-760), or pEGFP-N-OtDUB_GEF_-E572A-GFP with Lipofectamine 3000 (Invitrogen L3000001), and media was replaced 5 hours post-transfection.

### Microscopy

On the same day as fibroblast transfections, 12 mm coverslips were coated in 50 μg/ml poly-D-lysine (Corning Inc.) for 20 minutes at room temperature (RT), then washed 3 times with PBS. Coverslips were then coated in 10 μg/ml human fibronectin at 4°C overnight. The following day, coverslips were washed 3 times in PBS and blocked in 1% bovine serum albumin (BSA) for 1 h at 37°C. Coverslips were washed in PBS 3 more times before plating cells. 24 hours post transfection, 1.5×10^4^ cells were plated onto the coated coverslips.

After spreading on coated coverslips for 24 h (48 h after transfection), cells were fixed for 5 minutes in 2% paraformaldehyde in cytoskeleton buffer (10 mM MES pH 6.8, 138 mM KCl, 3 mM MgCl2, 2 mM EGTA, 320 mM sucrose). Cells were rinsed in TBS (150 mM NaCl, 20 mM Tris pH 7.4), permeabilized for 10 minutes in 0.3% Triton X-100/TBS, and then washed 3 times in 0.1% Triton X-100/TBS. They were then blocked for 30 minutes in Abdil (0.1% Triton X-100, 2% BSA, 0.1% NaAz, 10% FBS, TBS) and incubated with primary antibody (Abdil containing a 1:2000 dilution of Goat Anti-GFP (Rockland 600-101-215) at 4°C overnight.

The next morning, cells were washed in 0.1% Triton X-100/TBS 3 times and incubated in secondary antibody and phalloidin for 1 h at room temperature (RT) (in Abdil, 1:2000 Alexa Fluor 488 Donkey Anti-Goat (Abcam ab150129) and 1:250 Phalloidin Atto 647N (Sigma-Aldrich 65906). Cells were washed once in 0.1% Triton X-100/TBS, once in TBS, and then mounted onto glass slides using AquaMount (Lerner Laboratories 13800). After drying, coverslips were sealed using clear nail polish and imaged by epifluorescence on a microscope (model TE2000-S, Nikon) using a 20X objective, or with a 40x objective on a spinning disk confocal microscope (UltraVIEW VoX spinning disc confocal (Perkin Elmer), Nikon Ti-E-Eclipse).

After imaging cells, CellProfiler^44^ was used to detect and edit cell edges to compute all morphological metrics. 36-58 cells were measured per condition, and an ordinary one-way ANOVA with multiple comparisons was used to determine significant differences between groups.

#### Human cell lysate experiments

##### OtDUB:Rho GTPases co-IPs

HeLa cell pellets were collected 24 h after transfection and resuspended in lysis buffer (50 mM Tris-HCl pH 7.5, 150 mM NaCl, 0.2% Triton X-100, 2 mM PMSF, cOmplete protease inhibitor tablet (Roche)) and incubated on ice for 30 min with intermittent vortexing. The insoluble fraction was then pelleted by centrifuging at 21,000 x *g* for 15 min. Protein concentrations were determined by the Bradford assay (Bio-Rad) and normalized to 1 mg/ml with lysis buffer. IPs were carried out by rotating 850 μl lysate with15 μl of M2 anti-Flag resin (Sigma) at 4°C for 4 h. The resin was washed three times with 0.5 ml of lysis buffer, followed by protein elution at 37°C for 15 min with 50 μl of triple-Flag peptide elution buffer (0.4 mg/ml of peptide in lysis buffer). The eluent was isolated with a Quik-Spin Column (Bio-Rad), mixed with Laemmli sample buffer, boiled for 3 min and 25% was resolved by SDS-PAGE for immunoblotting.

##### Active Rac1/Cdc42 enrichment

At 32 h post-transfection, the HeLa cell medium was replaced with serum-free DMEM to reduce the basal levels of active Rac1/Cdc42 (PMID: 19948726). At 48 h post-transfection, cells were harvested and resuspended in Pak1PBD buffer (50 mM Tris-HCl pH 7.5, 150 mM NaCl, 0.2% Triton X-100, 5 mM MgCl_2_, 2 mM PMSF, cOmplete EDTA-free protease inhibitor tablet (Roche)). Lysates were clarified, quantified and normalized as above. Purified GST-Pak1PBD (20 μg) was added to 1 ml of 1.0 mg/ml lysates along with 15 μl of glutathione resin (ThermoFisher) and rotated for 90 min at 4°C. Resin was washed 3 times with 0.5 ml of Pak1PBD buffer and resuspended in Laemmli sample buffer. Samples were boiled for 3 min and 50% of eluted proteins were resolved by SDS-PAGE followed by immunoblotting.

### Immunoblotting

SDS-PAGE gels were transferred to PVDF (Millipore), blocked in 3% BSA in Tris-Buffered Saline containing 0.01% Tween-20 (TBST) or 5% dry milk in TBST. Blots were then incubated with primary antibodies diluted in TBST with 3% BSA or 5% milk for 1-2 h at RT or overnight at 4°C. Primary antibodies used in this study: mouse anti-Flag (F3165, Sigma), rabbit anti-Rac1/2/3 (2465, Cell Signaling), rabbit anti-Cdc42 (2466, Cell Signaling), rabbit anti-RhoA (2117, Cell Signaling), and rabbit anti-OtDUB (generated in-house). After washing, the filters were incubated with HRP-conjugated secondary antibodies in the respective blocking agent (NA931V and NA934V, GE Healthcare) for 1-2 h at RT or overnight at 4°C. Blots were visualized by enhanced chemiluminescence^45^, or SuperSignal West Pico PLUS (ThermoFisher) on a G:Box imaging system (Syngene). Densitometry was done in Gene Tools software (Syngene). Relative levels of active Rho GTPases were calculated by taking the active GTPase densitometry value divided by the input signal and normalized to the empty vector ratio for each experiment.

#### In vitro experiments

##### Purification of GST-tagged proteins

Rosetta DE3 cells transformed with pGEX-based plasmids were back-diluted in 2 L of LB supplemented with 100 μg/ml of ampicillin, grown at 37°C, and induced with 300 μM IPTG at OD_600_ 0.5-0.6. Induced cultures were then grown at 16-18°C for 16-19 h before harvesting. Pelleted *E. coli* cells were resuspended in ice-cold PBS containing 400 mM KCl, 1 mM DTT, 2 mM PMSF, lysozyme and DNase; incubated on ice for 1 h; lysed in a French press at 900 lbs.; and centrifuged at 50,000 x *g* for 60 min. Clarified lysates were incubated with 2 ml of glutathione resin and rotated for 1 h at 4°C in a 25 ml disposable drip column (Bio-Rad). Settled resin was washed with 25 column volumes (CV) of PBS + 400 mM KCl before eluting with 5-10 CV of 250 mM Tris-HCl pH 8, 0.5 M KCl, 10 mM reduced glutathione.

##### Purification of His-tagged OtDUB_GEF_, Rac1, and Cdc42

Transformed BL21(DE3) *E. coli* were grown in LB overnight, and back-diluted in Terrific Broth supplemented with 40 μg/ml kanamycin to OD_600_ 0.6 – 0.8 at 37°C, induced with 500 μM IPTG, and grown for 16 h at 18°C. Cells were harvested as above and resuspended in 50 mM Tris-HCl pH 8.0, 500 mM NaCl, 0.1 mM TCEP (Tris(2-carboxyethyl)phosphine) supplemented with cOmplete protease inhibitors (Roche). Clarified lysates were passed over 5 ml of packed Ni-NTA agarose resin (Qiagen) at 1.0 ml/min, washed with 15-20 CV of lysis buffer at 4 ml/min, and eluted with lysis buffer supplemented with 250 mM imidazole pH 8.0. Pooled elution fractions from Ni-NTA were concentrated in centrifugal filter units (Amicon; 10 K MWCO), and injected onto a HiLoad Superdex 75 pg column (GE Healthcare) equilibrated with 25 mM Tris-HCl pH 7.5, 100 mM NaCl, 0.1 mM TCEP. Proteins purified for crystallization were not cleaved of their affinity tags. Fractions containing protein complexes were identified by SDS-PAGE, pooled, concentrated to ∼50 mg/ml, flash frozen in liquid N_2_, and stored at −80°C.

##### GST-Pak1PBD

Eluates from glutathione affinity columns were concentrated and then resolved by FPLC (Akta) on a Superdex 75 Hiload 16/600 equilibrated with 50 mM TrisHCl pH 7.5, 150 mM NaCl. Peak fractions were pooled, concentrated and quantified by A260/280. The protein was diluted to 40 mg/ml, supplemented with MgCl_2_ to 5 mM final, flash frozen in 20 μl aliquots and stored at - 80°C. pGEXTK-Pak1 70-117 was a gift from Jonathan Chernoff (Addgene plasmid # 12217; http://n2t.net/addgene:12217; RRID:Addgene_12217)

##### GST, GST-Rac1, and Cdc42

Glutathione-resin eluates were dialyzed overnight in 4 L of 50 mM Tris-HCl, pH 7.5, 150 mM NaCl, 1mM DTT; concentrated; and quantified by A260/280. Aliquots were flash frozen and stored at −80°C.

##### GST-OtDUB_275-1369_-Flag and OtDUB_675-1369_-Flag

Glutathione-resin eluates were mixed with GST-HRV-3C protease and dialyzed overnight in 4 L PBS + 400 mM KCl, 1 mM DTT. Concentrated eluates were then resolved by FPLC on a Superose 6 column pre-equilibrated with 50 mM Tris-HCl, pH 7.5, 150 mM NaCl, 1mM DTT. Peak fractions were pooled. The OtDUB_675-1369_-Flag Superose 6 fractions required an additional GST depletion by incubating with glutathione resin prior to concentrating. Concentrated proteins were quantified by the BCA assay (ThermoFisher), and aliquots were flash frozen and stored at - 80°C.

##### Pulldown assays with GST-Rac1/Cdc42 and OtDUB_275-1369_-Flag/OtDUB_675-1369_-Flag

Protein combinations were diluted to 1 μM in binding buffer (50 mM Tris-HCl, pH 7.5, 150 mM NaCl, 0.01% Triton X-100) to a final volume of 500 μl. After incubating at 37°C for 30 min, the samples were spiked with 30 μl glutathione resin or M2-Flag resin and rotated for an additional 30 min at RT. Resin pellets were washed three times with binding buffer and eluted with Laemmli sample buffer or with 3xFlag peptide as described above.

##### In vitro pulldowns with GST-OtDUB fragments and Rac1

GST-tagged OtDUB fragments (20 μM) were mixed with Rac1 (60 µM) in binding buffer (50 mM Tris-HCl pH 7.5), 150 mM NaCl, 0.1 mM TCEP) to a final volume of 250 μL. Reactions were incubated with 200 μL glutathione resin at 4°C for 1 hour. The beads were washed with 15 CV binding buffer, and eluted with binding buffer supplemented with 10 mM reduced L-glutathione. Eluates were boiled and resolved by SDS-PAGE and stained with Coomassie Blue.

##### BODIPY-GDP nucleotide exchange assays

GTPases at a final concentration of 10 μM were loaded with sub-stoichiometric (1/4 [GTPase]) concentrations of GDP-BODIPY-FL (Thermo Scientific) in the presence of 50 mM Tris pH 7.5, 50 mM NaCl, 1 mM DTT, supplemented with 2 mM EDTA to promote unloading of co-purified nucleotide. Loading reactions were incubated for 1 hr at 25°C and then quenched by addition of MgCl_2_ to a final concentration of 25 mM. Exchange reactions were initiated by addition of a solution containing the OtGEF, 2 mM MgCl_2_, and 1 mM GTP. Reactions were mixed rapidly, loaded into a multi-well plate, and real-time fluorescence was collected on a plate reader for 30 mins at 25°C, using excitation at 488 nm and emission at 535 nm.^16^

##### Size exclusion chromatography (SEC)

Each OtDUB_GEF_ construct (75 μM) was mixed with the appropriate GTPase (150 μM) or diluted alone in a 500 μL reaction and incubated for 30 min at 4°C. Potential aggregates were removed by pelleting at 21,000 x *g* for 5 min prior to loading. Protein mixtures were injected into a Superdex 75 GL column (GE Healthcare) pre-equilibrated in 50 mM Tris pH 8.0, 100 mM NaCl, and 0.1 mM TCEP. Peak fractions were identified by monitoring at both 280 and 260 nm, resolved by SDS-PAGE, and visualized with Coomassie Blue staining.

##### Isothermal titration calorimetry (ITC)

The affinity of OtDUB_GEF_ for Rac1 and Cdc42 was determined using ITC with a NanoITC device (TA Instruments). All proteins were serially dialyzed four times against 25 mM HEPES pH 7.5, 50 mM NaCl for 12 hours each. Fifty μL of Rac1 (1.05 mM) was loaded into the syringe and injected into the cell containing 300 μL of OtDUB_GEF_ (150 μM) over 30 injections of 1.6 μL each. The resulting heats were measured at 25°C, and heats of dilution (buffer injected into OtDUB_GEF_) were subtracted. For Cdc42, the concentrations used were 3.15 mM of Cdc42 in the syringe, and 450 µM OtDUB_GEF_ in the cell. Data were collected in triplicate and analyzed using NanoAnalyze software; thermodynamic parameters were averaged over three runs.

#### Crystallography

##### Apo OtDUB_GEF_

Crystallization screening of apo OtDUB_GEF_ was performed by mixing the protein at ∼25 mg/mL in gel filtration buffer (25 mM Tris pH 7.0, 100 mM NaCl, 0.5 mM TCEP) with commercial crystallization formulations using the microbatch under-oil method at room temperature.^46^ Several crystallization hits were identified in a variety of conditions with the most promising one in 2.4 M sodium malonate pH 7.0. Optimizing ratio, pH, and temperature identified the final conditions. The final crystal was formed by streak seeding a drop composed of 2 μL of protein mixed with 2 μL of 2.4 M sodium malonate pH 7.0, mixed at room temperature and incubated underneath a ratio of 2:1 paraffin:silicone oil. Crystals were cryo-protected by gradual addition of crystallization buffer supplemented with 25% (v/v) glycerol and flash frozen in liquid N_2_. Diffraction data were collected at the Advanced Photon Source (APS) beamline 24-ID-C at 100 K temperature and wavelength 0.98 Å. The best crystal diffracted to 3.0 Å resolution.

Diffraction data were processed in HKL2000^47^ in space group *P*6_3_ and molecular replacement was performed using PHASER^48^ using the OtDUB_GEF_ molecule from the OtDUB_GEF_:Rac1 complex as a search model (see below). Six copies of the OtDUB_GEF_ were found in the asymmetric unit. The initial model was improved with iterative rounds of restrained refinement in REFMAC5^49^, and manual rebuilding in COOT^50^ using the 2Fo-Fc and Fo-Fc maps. Refinement statistics are summarized in Table 1.

##### OtDUB_GEF_:Rac1 complex

Crystallization screening of the OtDUB_GEF_:Rac1 complex was performed by concentrating the protein at ∼50 mg/mL in GEF buffer (25 mM Tris pH 7.0, 100 mM NaCl, 0.5 mM TCEP, 5 mM EDTA) at 25°C with commercial crystallization formulations using the microbatch under-oil method at RT. Several crystallization hits were identified after 24 hours; plate-shaped crystals formed in conditions with slightly basic pH (e.g. CHES pH 9.5, Tris-HCl pH 8.5) and moderate concentrations of PEG (20-25% PEG 3350 and 8000). pH vs. PEG grid screening revealed optimized conditions that formed thicker, single plate crystals. The final crystal was obtained by introducing microseeds^51^ to a drop composed of 2 μL of protein mixed with 2 μL of 25 mM CHES pH 9.5, 23% (v/v) PEG 8000. Crystals were cryo-protected by gradual addition of crystallization buffer supplemented with 25% (v/v) glycerol and flash frozen in liquid N_2_. Diffraction data were collected at the APS beamline 24-ID-C at 100 K temperature and wavelength 0.98 Å. The best crystal diffracted to 1.7 Å resolution.

Diffraction data were processed in HKL2000^47^ in space group *P*1 and molecular replacement was performed with PHASER^48^ using a monomer of unliganded Rac1 from the Tiam1:Rac1 structure (PDB: 1FOE) as a search model. Two copies of Rac1 were readily found within the asymmetric unit, and refined in REFMAC5.^49^ The high resolution of the diffraction data allowed for the automatic model building of OtDUB_GEF_ using the BUCCANEER^52^ autobuild function together with iterative rounds of refinement in REFMAC5 (keywords “ncyc 40”, “RIDG DIST SIGM 0.04”, “NCSR LOCAL”, “solvent YES”). After several rounds of automated mainchain tracing, more than 95% of the OtDUB_GEF_ had been built with good statistics. In total, the unit cell contained two copies of Rac1 and two copies of OtDUB_GEF_. The final model was produced after iterative rounds of restrained refinement in REFMAC5 and manual rebuilding in COOT^50^ using the 2Fo-Fc and Fo-Fc maps. Refinement statistics are summarized in Table 1.

##### Mass spectrometry sample preparation

GST, GST-OtDUB_275-1369_-Flag, and OtDUB_675-1369_-Flag were processed as above from 1L LB cultures except instead of eluting the polypeptides from the resin, the resin-bound proteins were incubated with HeLa cell lysates. The HeLa cell lysates were generated by scraping off and pooling 17 confluent 15-cm plates. The resulting cell pellet was lysed in 30 ml of lysis buffer by vortexing and rotating for 20 min at 4°C (lysis buffer: 50 mM Tris-HCL pH 7.5, 150 mM NaCl, 0.5% Triton X-100, 1 mM DTT, 2 mM PMSF, cOmplete protease inhibitor tablet). The insoluble fraction was removed by centrifugation at 30,000 x *g* for 20 min at 4°C, and the clarified lysates (∼2 mg/ml) were split equally across the three columns containing GST, GST-OtDUB_275-1369_-Flag, or OtDUB_675-1369_-Flag. The columns were rotated for 2 h at 4°C, and the settled resins were washed with 33 CV of wash buffer (50 mM Tris-HCL pH 7.5, 150 mM NaCl), then eluted by an overnight on-column cleavage with GST-HRV3C in wash buffer. Elutes were concentrated with a 3500 MW cutoff Amicon Ultra filter (Millipore) and submitted as in-solution samples to the Keck Mass Spectrometry Facility at Yale University. Duplicates of each identical sample were submitted to the facility for independent processing.

##### In-solution protein digestion

Proteins were precipitated from the eluates with acetone using established protocols.^53^ Protein pellets were dissolved and denatured in 8 M urea, 0.4 M ammonium bicarbonate, pH 8. The proteins were reduced by the addition of 1/10 volume of 45 mM dithiothreitol (Pierce Thermo Scientific #20290) and incubation at 37°C for 30 min, then alkylated with the addition of 1/10 volume of 100 mM iodoacetamide (Sigma-Aldrich #I1149) with incubation in the dark at RT for 30 min. The urea concentration was adjusted to 2M by the addition of water prior to enzymatic digestion at 37°C with trypsin (Promega Seq. Grade Mod. Trypsin, # V5113) for 16 h. Protease:protein ratios were estimated at 1:50. Samples were acidified by the addition of 1/40 volume of 20% trifluoroacetic acid, then desalted using C18 MacroSpin columns (The Nest Group, #SMM SS18V) following the manufacturer’s directions with peptides eluted with 0.1% TFA, 80% acetonitrile. Eluted peptides were dried in a Speedvac, dissolved in MS loading buffer (2% aceotonitrile, 0.2% trifluoroacetic acid in water) and their concentrations determined (A260/A280; Thermo Scientific Nanodrop 2000 UV-Vis Spectrophotometer). Each sample was then further diluted with MS loading buffer to 0.05 µg/µl, with 0.25 µg (5 µl) injected for LC-MS/MS analysis.

##### LC-MS/MS

Analysis was performed on a Thermo Scientific Q Exactive Plus equipped with a Waters nanoAcquity UPLC system utilizing a binary solvent system (A: 100% water, 0.1% formic acid; B: 100% acetonitrile, 0.1% formic acid). Trapping was performed at 5 µl/min, 99.5% Buffer A for 3 min using a Waters Symmetry® C18 180 µm x 20 mm trap column. Peptides were separated using an ACQUITY UPLC Peptide BEH C18 nanoACQUITY Column 1.7 µm, 75 µm x 250 mm (37°C) and eluted at 300 nl/min with the following gradient: 3% buffer B at initial conditions; 5% B at 1 min; 25% B at 90 min; 50% B at 110 min; 90% B at 115 min; 90% B at 120 min; return to initial conditions at 125 min. MS was acquired in profile mode over the 300-1,700 m/z range using 1 microscan, 70,000 resolution, AGC target of 3E6, and a maximum injection time of 45 ms. Data dependent MS/MS were acquired in centroid mode on the top 20 precursors per MS scan using 1 microscan, 17,500 resolution, AGC target of 1E5, maximum injection time of 100 ms, and an isolation window of 1.7 m/z. Precursors were fragmented by HCD activation with a collision energy of 28%. MS/MS were collected on species with an intensity threshold of 1E4, charge states 2-6, and peptide match preferred. Dynamic exclusion was set to 20 seconds.

##### Peptide identification

Data were analyzed using Proteome Discoverer software v2.2 (Thermo Scientific). Data searching was performed using the Mascot algorithm (version 2.6.1) (Matrix Science) against a custom database containing protein sequences for the OTT_1962 constructs of interest as well as *Escherichia coli* and human proteomes (derived from the Swissprotein database). The search parameters included tryptic digestion with up to 2 missed cleavages, 10 ppm precursor mass tolerance and 0.02 Da fragment mass tolerance, and variable (dynamic) modifications of methionine oxidation and carbamidomethylated cysteine. Normal and decoy database searches were run, with the confidence level set to 95% (p<0.05). Scaffold (version Scaffold_4.10.0, Proteome Software Inc., Portland, OR) was used to validate MS/MS based peptide and protein identifications. Peptide identifications were accepted if they could be established at greater than 95.0% probability by the Scaffold Local FDR algorithm. Protein identifications were accepted if they could be established at greater than 99.0% probability and contained at least 2 identified peptides. Peptide counts from each independent mass spectrometry processing were then added together for each sample. The full dataset is available as Supplementary Table 1.

##### Accession numbers

Coordinates and structure factors have been deposited in the Protein Data Bank with PDB ID: 6×1H (OtDUB_GEF_ apo) and 6×1G (OtDUB_GEF_:Rac1 complex).

## Supplemental Figure Legends

**Figure S1.**
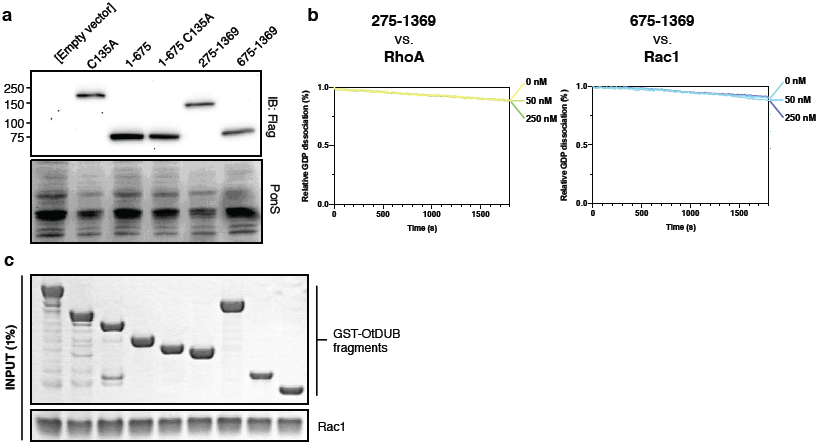
Expression, activity, and catalytic selectivity of OtDUB fragments. (a) OtDUB fragments expressed in yeast. Transformants were grown for 4 h in minimal liquid medium containing galactose. After induction, equivalent optical density units (ODU) were processed and extracts from 0.25 ODU were resolved by SDS-PAGE and immunoblotted with anti-Flag antibody, then Ponceau stained (PonS) for loading comparison. Absorbance by dead cells results in reduced Ponceau signal for C135A and 275-1369. (b) Fluorescent nucleotide exchange assays demonstrate absence of activity against RhoA in the OtDUB_275-1369_ fragment (left), and absence of activity against Rac1 in truncation construct OtDUB_657-1369_. (c) Input proteins from pulldown experiment in Figure 3A.

**Figure S2.**
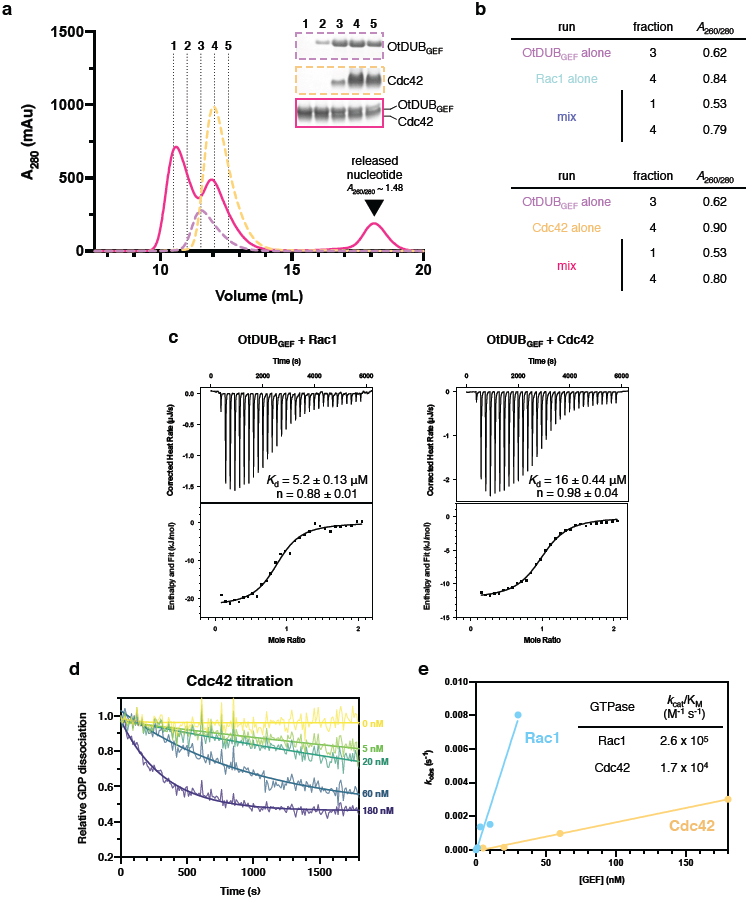
Binding studies of OtDUB_GEF_ and Rac1 or Cdc42. (a) Size exclusion chromatography (SEC) of OtDUB_GEF_:Cdc42 mixtures, as in Figure 3B. (b) *A*_260/280_ ratios for fractions in each SEC run. OtDUB_GEF_ alone and GTPase alone runs have *A*_260/280_ of 0.6-0.9, whereas complex peaks formed after mixing are devoid of nucleotide (*A*_260/280_∼ 0.5). Exchanged nucleotide is also observed at late elution volumes (triangle). (c) Isothermal titration calorimetry quantification of binding affinity for OtDUB_GEF_ and Rac1 (K_d_ ∼ 5 μM) or Cdc42 (K_d_ ∼ 16 μM). (d) Fluorescent nucleotide exchange assays as in Figure 3C, except with Cdc42 instead of Rac1, with increasing concentrations of OtDUB_GEF_. (e) Linear transformation and calculated *k*_cat_/*K*_M_ values for Rac1 (2.6 ± 0.3 ×10^5^ M^-1^ s^-1^) and Cdc42 (1.7 ± 0.1 ×10^4^ M^-1^ s^-1^).

**Figure S3.**
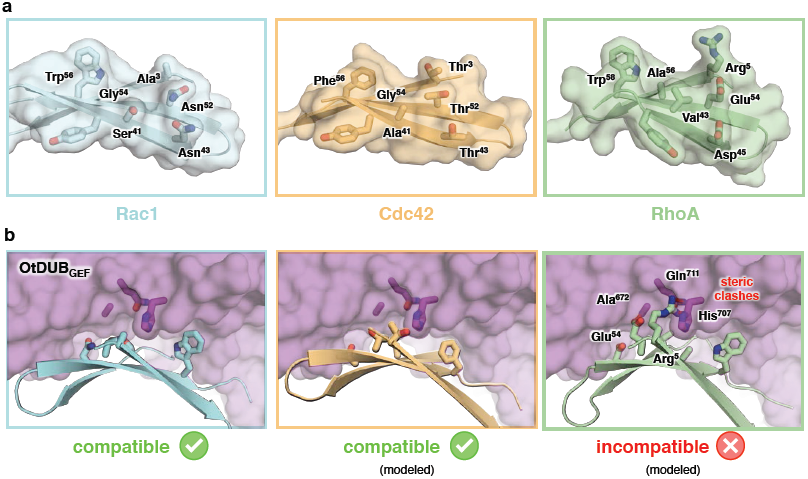
Structural basis for GTPase selectivity. (a) β2-3 hairpin “interswitch” regions from Rac1, Cdc42, and RhoA are shown as cartoon and transparent surface, with surface-exposed residues as sticks. (b) Structural alignment of the selectivity patches of Cdc42 and RhoA reveal steric clashes with RhoA that likely prevent binding with OtDUB_GEF_ (purple surface and sticks)

**Figure S4.**
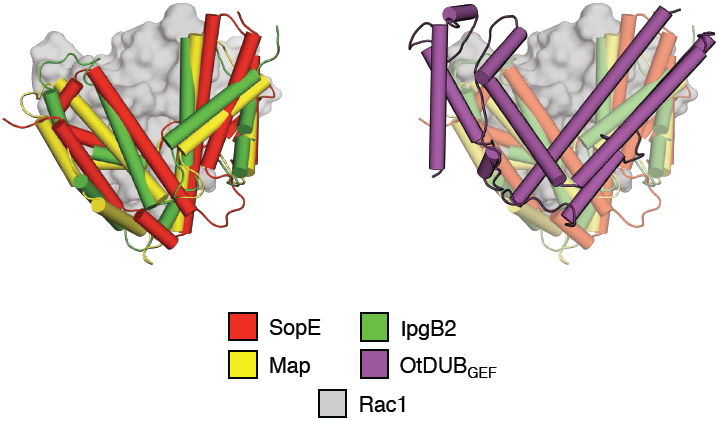
OtDUB_GEF_ occupies a similar footprint on Rac1 as other bacGEFs. Structural alignment by GTPase (surface) of available bacGEFs (cylinders): SopE (red), Map (yellow), IpgB2 (green), and OtDUB_GEF_ (purple) highlight convergence on a conserved V-shaped fold on the same location on Rac1.

**Figure S5.**
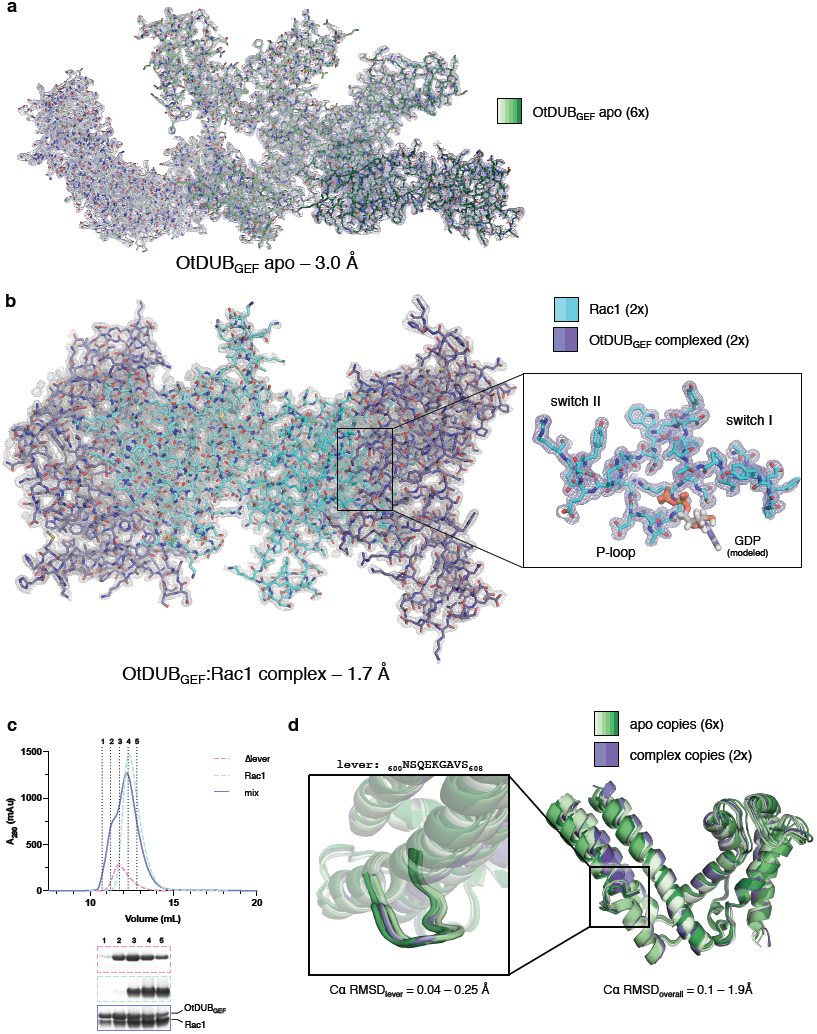
Overall structures of OtDUB_GEF_ apo and Rac1 complexes. (a) Overall crystal structure of OtDUB_GEF_ resolved to 3.0 Å, with six copies in the asymmetric unit (ASU), with the 2Fo-Fc electron density map (mesh) contoured at 1.0 σ. Pairwise Cα RMSDs between copies in the ASU ranges from 0.36 to 1.9 Å. (b) Overall crystal structure of OtDUB_GEF_ in complex with Rac1 resolved to 1.7 Å, with the 2Fo-Fc electron density map (mesh) contoured at 1.0 σ. The asymmetric unit contains two copies each of OtDUB_GEF_ and of Rac1. Each complex is virtually identical, with Cα RMSD between complexes of 0.1 Å. Inset: detailed view of the nucleotide-binding region of Rac1 reveals absence of electron density for GDP (modeled, grey). (c) Size exclusion chromatography of OtDUB_GEF/Δlever_:Rac1 mixtures. The late eluting peak representing exchanged nucleotide, which was observed with wild-type OtDUB_GEF_, is absent. (d) Structural comparison of six copies of OtDUB_GEF_ from the apo structure (green shades) and two copies of OtDUB_GEF_ from the Rac1 complex structure (purple shades). Inset: detailed view of the catalytic lever region reveals a rigid conformation in both apo and complex OtDUB_GEF_ molecules.

**Figure S6.**
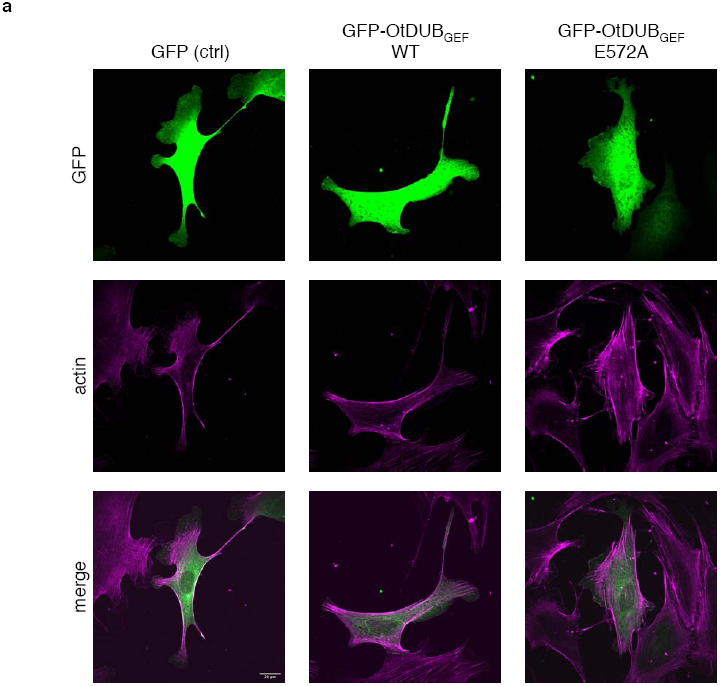
Confocal microscopy of fibroblasts expressing OtDUB_GEF_. Representative immunofluorescence images of cells transfected with plasmids expressing GFP, GFP-OtDUB_GEF_ wild-type (WT) or E572A mutant. No gross changes are observed in actin cytoskeleton structure, as visualized with phalloidin-Atto-N647N staining (purple).

**Figure S7.**
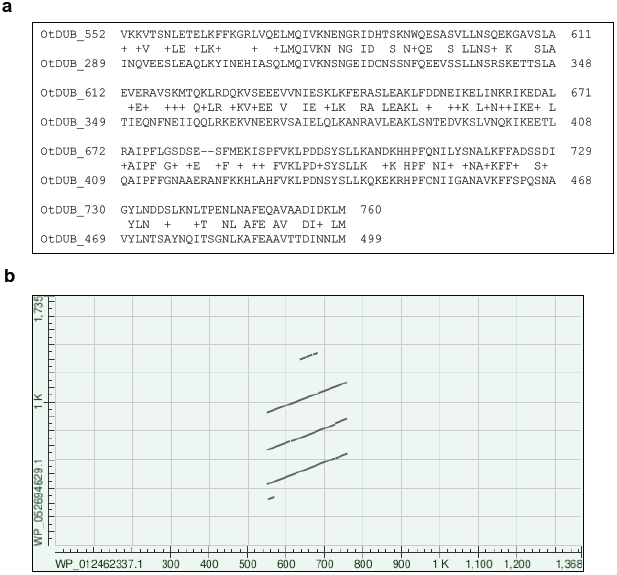
A second OtDUB_GEF_-related sequence is present in OtDUB. (a) Sequence alignment using BLASTP revealing a high-identity duplication of the OtDUB_GEF_ within residues 289-499. (b) A pairwise alignment of OtDUB (WP_012462337.1) residues 548-759 against full-length OcDUB (WP_052694629.1) within BLASTp reveals three high identity repeats within OcDUB. E572 is conserved in two of the three sequences in OcDUB: range 1 (928-1137) and range 2 (429-641), but not within range 3 (668-883).

**Figure S8.**
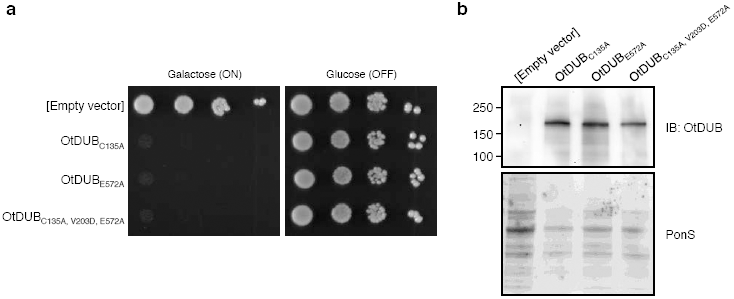
A FL OtDUB derivative lacking GEF activity is still toxic in yeast. (a) Growth assay of W303 yeast transformed with p416GAL1 plasmids expressing various OtDUB mutants in the presence of galactose. This OtDUB mutant also lacked deubiquitylase and strong ubiquitin-binding activities. Yeast cultures were serially diluted in ten-fold steps and spotted on SD-URA plates containing either galactose or glucose as tbe carbon source and grown for 3 days at 30°C. (b) Expression of OtDUB protein fragments in yeast. Transformants were induced for expression by growing for 4 h in minimal media containing galactose. After induction, equivalent ODUs were processed, and 0.25 ODU were resolved by SDS-PAGE and immunoblotted with an anti-OtDUB antibody, followed by Ponceau S staining (PonS) to compare loading. Reduced Ponceau signal in OtDUB expressing strains is due to absorbance by dead yeast.

